# From computing transition probabilities to word recognition in sleeping neonates, a two-step neural tale

**DOI:** 10.1101/2021.07.16.452631

**Authors:** Ana Fló, Lucas Benjamin, Marie Palu, Ghislaine Dehaene-Lambertz

## Abstract

Extracting statistical regularities from the environment is a primary learning mechanism, which might support language acquisition. While it is known that infants are sensitive to transition probabilities between syllables in continuous speech, the format of the encoded representation remains unknown. Here we used electrophysiology to investigate how 31 full-term neonates process an artificial language build by the random concatenation of four pseudo-words and which information they retain. We used neural entrainment as a marker of the regularities the brain is tracking in the stream during learning. Then, we compared the evoked-related potentials (ERP) to different triplets to further explore the format of the information kept in memory. After only two minutes of familiarization with the artificial language, we observed significant neural entrainment at the word rate over left temporal electrodes compared to a random stream, demonstrating that sleeping neonates automatically and rapidly extracted the word pattern. ERPs significantly differed between triplets starting or not with the correct first syllable in the test phase, but no difference was associated with later violations in transition probabilities, revealing a change in the representation format between segmentation and memory processes. If the transition probabilities were used to segment the stream, the retained representation relied on syllables’ ordinal position, but still without a complete representation of the words at this age. Our results revealed a two-step learning strategy, probably involving different brain regions.

## 1. Introduction

From before birth, infants demonstrate learning capacities. During the last weeks of gestation, they learned some prosodic features of their native language (Mehler et al., 1988) and their mother’s voice (DeCasper & Fifer, 1980), as the taste of the amniotic liquid (Marlier et al., 1998), and rapidly learn to recognize their mother’s face after birth (Bushneil et al., 1989). Neonates also quickly adapt to repeated sensory information. For example, after a few minutes of familiarization with a word, they notice when it changed (Benavides-Varela et al., 2012; Benavides-Varela & Mehler, 2011), and similar short-term memories have been described for previously seen faces (Pascalis et al., 1995, p. 994). Despite these undeniable learning and memory capacities, very little is known about the underlying mechanisms, the information neonates are sensitive to, and the format of the representation in which information is memorized.

Here we focused on a primary yet indispensable fast learning mechanism: statistical learning. Statistical learning refers to the capacity to detect regularities in the input. Abundant literature (Saffran & Kirkham, 2018) shows that this mechanism is common across domains (visual, auditory) (Bulf et al., 2011; Fiser & Aslin, 2002; Kirkham et al., 2002; Saffran et al., 1996, 1999), species (primates, rodents) (Hauser et al., 2001; Toro & Trobalón, 2005), and extends to different levels of stimulus/scene complexity. Concerning language acquisition, statistical learning has been proposed as a critical mechanism to explain how infants might discover linguistic regularities. For example, it might serve to identify word candidates based on frequently co-occurring syllables (Saffran et al., 1996), to discover phonotactic and acoustic patterns (Friederici et al., 2007; Jusczyk et al., 1999), and to detect morphological and syntactic regularities (Shi et al., 1999).

Experimental evidence supporting the role of statistical learning in language acquisition has been mainly obtained in word segmentation tasks from an artificial speech stream in which acoustic cues have been removed. In a seminal study (Saffran et al., 1996, p. 996), 8-month-old infants were first exposed to 3 minutes of an artificial speech (thereafter called Structured stream) constituted by four randomly concatenated tri-syllabic pseudo-words, with the drops of transition probabilities (TPs) between syllables as the only cue to word boundaries. Within a pseudo-word, the first two syllables predict the following syllable (TP equal to 1), while the last syllable could be followed by any other pseudo-words (TP equal to 1/3). When test triplets were then played in isolation, infants’ looking pattern differed between the pseudo-words (i.e., Words: both TPs in the triplet equal 1) and triplets straddling a TP drop (i.e., Part-words: one TP equal 1 and the other equal 1/3). This result, uncovered that infants are sensitive to the statistical relations between syllables, yet, it remains unknown what they exactly learn.

It is commonly assumed that infants segment the stream into words that are memorized and subsequently recognized when presented in isolation. However, two other hypotheses can also explain the results. Infants may compute the transitional probabilities matrix between all syllables through synaptic plasticity and Hebbian learning (Endress & Johnson, 2021) without segmenting the stream (Benjamin et al., in prep.). The different association strength between syllables in Words and Part-Words could support the difference between these conditions. Alternatively, infants may segment the stream using the drop of transitional probabilities at the end of the Words but only memorize the syllable following the drop. Indeed, because this syllable being less predictable during the stream, it might induce surprise, a powerful learning factor for infants (Stahl & Feigenson, 2015). The three hypotheses are not dissociable in the existing studies since they all result in differential responses to Words and Part-words. Nevertheless, each explanation relies on different mechanisms in terms of computational complexity and neural bases.

A crucial difference between encoding the TPs matrix and segmenting the stream into Words is that memory constraints for sequence encoding may enter into play. When encoding a sequence of items, each item is associated with the close items (i.e., TPs) and its ordinal position within the sequence (Henson, 1998). Indeed, Dehaene et al. (Dehaene et al., 2015) proposed a taxonomy of five levels along which a sequence can be encoded: from *(1)* TPs between elements, passing by *(2)* chunking (grouping of elements in a unit), and *(3)* ordinal knowledge (the elements have an ordered position in the unit), until more abstract encoding based on *(4)* rules and *(5)* nested structures. In a very recent study in 23 adult patients with implanted electrodes who listened to a structured stream containing Words, the first stages of this taxonomy were explored using representational similarity analyses. The authors reported a complex picture in which different brain regions hosted different representations (Henin et al., 2021). Some electrodes, located in the superior temporal gyrus and the pars opercularis and motor cortex, were responding to TPs encoding. Others, located in the inferior frontal gyrus, anterior temporal lobe, and posterior superior temporal sulcus, were sensitive to ordinal position (first vs. second vs. third syllable). Finally, in the hippocampus, electrodes were sensitive to Words (chunks). Given the complex maturational calendar of the different brain structures, particularly the slow maturation of the hippocampus (Lavenex & Banta Lavenex, 2013) and frontal areas (Lebenberg et al., 2019), the generalization of these results to young infants might be inadequate. Furthermore, attention is notably limited at a young age, especially in neonates who sleep most of the time. Thus, we may wonder whether passive exposure might be sufficient or whether some of these computations, such as representing syllables’ ordinal position and active prediction of the next item, might not be observed during sleep. In other words, our goal was to study which levels of this taxonomy newborns possess to support language acquisition.

Three previous studies have shown that neonates are at least sensitive to the first level, TPs encoding. During a long familiarization with an artificial stream of syllables (15 mn) (Teinonen et al., 2009) and of tones (9 mn) (Kudo et al., 2011), a different event-related responses emerged to the first syllables/tones of the Words. However, this result may reflect either the response to a local prediction error (i.e., TPs), or to changes in the syllable frequencies as the medial and last syllable were repeated from time to time, or to truly individual triplets. A third study using Near-Infrared Spectroscopy (NIRS) showed a differential BOLD response to Words and Part-words following a 3.5 minutes familiarization with the structure stream (Fló et al., 2019). While adding that neonates can remember the extracted information for a few minutes, it leaves pending the information they retained that triggered the differential response. We, therefore, propose here to further investigate the study of statistical learning in neonates using high-density EEG (128 electrodes) in a paradigm based on three minutes of exposure to a structured stream, followed by the presentation of isolated triplets in a test phase (Fig 1). Our goal was double, first to target learning cues during the stream exposure thanks to neural entrainment, and second to characterize the format of the learned representation by presenting four different types of triplets in isolation.

**Figure 1.**
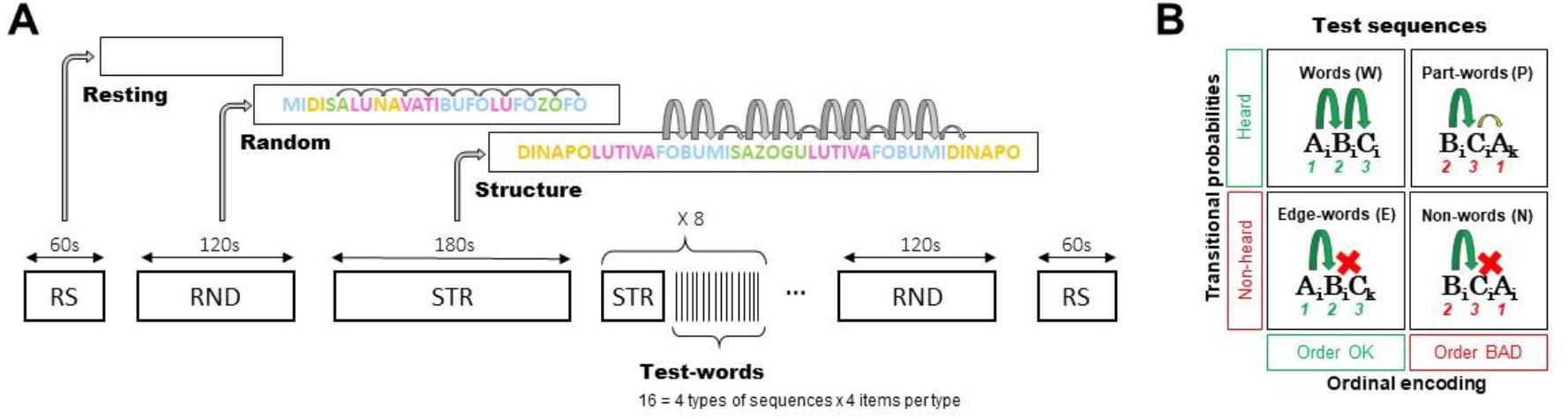
**(A)** Experimental protocol. A long Structured stream (180 s) was followed by 8 test short familiarizations (30 s) and test blocks, where 16 triplets were presented in isolation (ISI 2-2.5 s). Resting state and a Random stream were sandwiching the entire experiment to control for the effect of time, and notably infant vigilance, on the EEG recordings. **(B)** Possible types of test words. Test words could have violations in the TPs between the second and third syllables, in the ordinal position of the syllables, or both.

Thanks to its temporal sensitivity, EEG allows monitoring learning, even in non-participating subjects, such as sleeping neonates. In particular, in this paradigm, where syllables have a fixed duration, the auditory response induced by the regular presentation can be captured as entrainment at the frequency of stimulation (f=1/syllable duration). Crucially, this steady-state response is not limited to low-level features like syllable onset but can reflect any regular pattern the brain is tracking. Thus, if the listener detects the 3-syllabic pattern embedded in the stream, entrainment should also be observed at the triplet frequency (1/3 of the syllabic rate) (Batterink & Choi, 2021; Benjamin et al., 2021; Buiatti et al., 2009; Kabdebon et al., 2015). Here we quantified the entrained neural responses at the syllabic and Word rates measuring an enhanced Power and Inter Trial Coherence (ITC) during the presentation of the Structured stream, a Random stream (random concatenation of the syllables), and Resting-state (i.e., no stimulation). We expected a similar entrainment at the syllabic rate for both streams relative to resting-state, but an increased activity at the word rate during the Structured stream.

However, neural entrainment at the word frequency can result from two different processes, similarly to differential ERPs to syllables during the stream (Kudo et al., 2011; Teinonen et al., 2009). Either the neonates have detected a drop in TPs, or they recognize the triplets re-occurrence. To test what they learned and memorized, we compared the ERPs to isolated triplets in a post-learning phase (Fig 1B). We build four types of triplets to disentangle different hypotheses on the encoding format of the memorized pattern. We contrasted: *(1)* triplets respecting, or not, TPs between syllables, and *(2)* triplets violating, or not, the ordinal position of the first syllable. Therefore, we presented the classical conditions: Words (*A_i_B_i_C_i_*) corresponding to the pseudo-words present in the stream, and Part-words (*B_i_C_i_A_k_*) corresponding to triplets straddling a TP drop. Note that Part-words also have an incorrect first syllable. To these common conditions, we added two other conditions: Edge-words and Non-words. Edge-words (*A_i_B_i_C_k_*) were triplets in which the last syllable between two Words was exchanged; thus, they retained the ordinal position of the syllables, but they were never presented in the stream (last TP equaled zero). Non-words (*B_i_C_i_A_i_*) were triplets in which the first syllable appeared in the last position; thus, all syllables belonged to the same Word, but the ordinal position was incorrect, and the triplet was never heard (last TP equaled zero).

If neonates segment the stream and encode ordinal information or at least the first syllable of a word, we expected an early differential response between *ABx* (Words and Edge-Words) and *BCx* triplets (Part-Word and Non-Words). Note that any difference before the third syllable can only be due to the encoding of the first syllable —*A_i_B_i_* and *B_i_C_i_* had both TPs equal to one. By contrast, if the response to the isolated triplets only depends on the adherence to the statistical structure of the Structured stream, the ERPs between never heard triplets (Edge-words and Non-words) and those present in the stream (Words and Part-words) should differ from the third syllable. For the sake of completeness, we also considered that memory encoding following segmentation might be sensitive to the temporal proximity of the elements belonging to the same chunk as a community structure, predicting that Non-Words (*B_i_C_i_A_i_*) are closer to Words (*A_i_B_i_C_i_*) than Part-Words (*B_i_C_i_A_k_*).

Additionally, we tested 32 adult participants in a behavioral online experiment analog to the infant task. After familiarization with the structured stream, participants had to rate their familiarity with the test triplets. Because the stimuli (duration of the Structured streams and number of tests words) were the same as in the neonates’ study, this experiment provides a reference point of what mature and expert participants encode and memorize.

To summarize, simple TP learning should result in a difference between triplets present or not in the stream (Words+Part-words vs. Edge-words+Non-words). Segmentation should be revealed by neural entrainment at the word rate during the Structured stream and a difference between *ABx* and *BCx* sequences. The granularity of the encoding can be further investigated by comparing Words vs. Edge-Words and Non-words vs. Part-Words.

## 2. Materials and Methods

### 2.1. Participants

Participants were healthy-full-term neonates, with normal pregnancy and birth (GA > 38 weeks, Apgar scores ≥ 7/8 in the 1/5 minute, birthweight > 2.5 Kg, cranial perimeter ≥ 33.0 cm), tested at the Port Royal Maternity (AP-HP), in Paris, France. Parents provided informed consent. 31 participants who provided enough data without motion artifacts were included (10 females; 1 to 4 days old; mean GA: 40.2 weeks; mean weight: 3475 g). Seven other infants were excluded from the analysis (3 due to excessive hair or cradle cap, 2 due to excessive motion artifacts, and 2 because the parents decided to interrupt the experiment).

### 2.2. Stimuli

The stimuli were synthesized using the fr4 French female voice of the MBROLA diphone database (Dutoit et al., 1996). Syllables had a consonant-vowel structure. Each phone had a duration of 125 ms and a constant pitch of 200 Hz. The streams were continuous with coarticulation and no pauses, and they were ramped up and down during the first and last 5 s to avoid the start and end might serve as perceptual anchors. The structured streams consisted of a semi-random concatenation of the four tri-syllabic pseudo-words. Pseudo-words were concatenated with the only restrictions that the same word could not appear twice in a row, and the same two words could not repeatedly alternate more than two times (i.e., the sequence *W_k_W_j_W_k_W_j_*, where *W_k_* and *W_j_* are two words, was forbidden). The learning stream lasted 180 seconds, each word appearing 60 times and each of the 12 possible part-words 18 to 21 times; the average TPs between words was 0.332 (SD = 0.017, range 0.310 to 0.361). The eight short structured streams lasted 30 seconds each, each word appearing 80 (8×10) times and each of the 12 possible part-words between 24 and 28 times; the average transitional probability between words was 0.325 (SD = 0.012, range 0.308 to 0.345). The pseudo-words were created to avoid that specific phonetic features could help to segment the stream. Additionally, three different structured streams (lists A, B, and C, corresponding to three sets of pseudo-words) were used by modifying how the syllables were combined to form the Words (Table 1). Participants were randomly assigned and balanced between lists.

**Table 1.** Triplets for each condition (Words, Edge-words, Part-words, and Non-words) used for each of the three lists.

| List | A<sub>i</sub>B<sub>j</sub>C<sub>i</sub> | A<sub>i</sub>B<sub>j</sub>C<sub>k</sub> | B<sub>i</sub>C<sub>i</sub>A<sub>k</sub> | B<sub>i</sub>C<sub>i</sub>A<sub>i</sub> |
| --- | --- | --- | --- | --- |
| A | dinapo | dinava | napolu | napodi |
|  | lutiva | lutimi | tivafo | tivalu |
|  | fobumi | fobugu | bumisa | bumifo |
|  | sazogu | sazopo | zogudi | zogusa |
| B | napolu | napofo | poluti | poluna |
|  | tivafo | tivasa | vafobu | vafoti |
|  | bumisa | bumidi | misazo | misabu |
|  | zogudi | zogulu | gudina | gudizo |
| C | poluti | polubu | lutiva | lutipo |
|  | vafobu | vafozo | fobumi | fobuma |
|  | misazo | misana | sazogu | sazomi |
|  | gudina | guditi | dinapo | dinagu |

The random were built a semi-random concatenation of the same 12 syllables, with the only restriction that the same syllable could not appear twice in a row and that two syllables could not alternate more than two times (i.e., the sequence *S_k_S_j_S_k_S_j_*, where *S_k_* and *S_j_* are two syllables, was forbidden). Streams lasted 120 seconds each, and each syllable appeared 480 times in each of them.

Test words were tri-syllabic triplets presented in isolation.

### 2.3. Procedure and data acquisition

Scalp electrophysiological activity was recorded using a 128-electrode net (Electrical Geodesics, Inc.) referred to the vertex with a sampling frequency of 250 Hz. Neonates were tested in a soundproof booth while sleeping or during quiet rest. The study involved: *(1)* 60 s of resting-state; *(2)* 120 s of a random stream; *(3)* 180 s of a structured stream *(4)* 8 series of a 30 s of structured streams followed by 16 test sequences (ISI 2-2.5s); *(5)* 120 s of a random stream; *(6)* 60 s of resting state. The random streams and resting-state were presented before and after the learning and test parts to avoid sensing entrainment differences due to time in the experiment (i.e., a change in the vigilance state).

### 2.4. Data pre-processing

Data were band-pass filter 0.1-40 Hz and pre-processed using custom MATLAB scripts based on the EEGLAB toolbox 2021.0 (Delorme & Makeig, 2004), according to the APICE pre-processing pipeline (Fló et al., 2021). The following steps were applied: *(1)* Data was band-pass filter 0.1-40 Hz. *(2)* Artifacts were identified based on voltage amplitude, variance, first derivative, and running average using adaptive thresholds. *(3)* Time samples were defined as artifacts if more than 30% of the channels presented artifacts. *(4)* Electrodes were spatially interpolated using spherical splines for samples that were not marked as bad.

### 2.5. Neural entrainment

The pre-processed data were resampled to 300 Hz to achieve an integer number of samples per triplet (225 samples in 0.75 s) and further high-pass filtered at 0.2 Hz. Then, data was segmented from the beginning of each phase into 0.75 s long segments. Segments containing samples with artifacts were rejected. On average we retained 74 % of the data during Resting (SD 17, range [31, 100]), 84 % of the data during the Random (SD, 11, [47, 100]), and 87 % of the data during the Structured (SD 7, range [71, 100]). Subjects who did not provide at least 6 segments per condition were not included.

#### Neural entrainment per condition

The 0.75 s epochs belonging to the same condition were reshaped into non-overlapping epochs of 7.5 s (10 triplets, 30 syllables), retaining the chronological order; thus, the timing of the steady state response. Data were referenced average and normalized by dividing by the standard deviation within an epoch. DSS, a technique based on spatial filters designed to remove stimulus-unrelated activity (de Cheveigné & Simon, 2008), was applied, and the first 30 components of the first PCA and the first 6 of the DSS filter were retained (the pattern of results did not differ if DSS was not used). Then, data were converted to the frequency domain using the Fast Fourier Transform (FFT) algorithm, and the power and ITC were estimated for each electrode during each condition (Resting-state, Random, Structured). Then, the SNR relative to the twelve adjacent frequency bins (six of each side corresponding to 0.8 Hz) was estimated. The power was computed as the power spectrum of the average response across trials. Then, the noise level was estimated at each frequency by assuming a power-law fit on the adjacent frequency bins *log(P_estimate_(f))* = *a*+*b*\**log(f)*. The SNR for the power was *SNR(f)* = *(log(P(f))-mean(P_noise_(f)))*/ *std(P_noise_(f))*, where *P_noise_(f)* = *log(P_estimate_(f))-log(P)*. The ITC was computed as 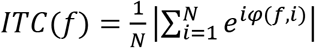, where *N* is the number of trials and *φ(f,i)* is the phase at frequency *f* and trial *i*. The ITC ranges from 0 to 1 (i.e., completely desynchronized activity to perfectly phased locked activity). The SNR was also estimated using the twelve adjacent frequency bins (six of each side corresponding to 0.8 Hz) as *SNR(f)* = *(ITC(f)-mean(ITC_noise_(f)))*/*std(ITCnoise(f))*, where *ITC_noise_(f)* is the ITC over the adjacent frequency bins.

For statistical analysis, the SNR was compared at the syllabic rate (4 Hz) and word rate (1.33 Hz) against zero (null hypothesis) using a one-tail t-test. P-values were corrected across electrodes by FDR.

#### Neural entrainment time course

The 0.75 s epochs were concatenated chronologically (1 minute of RS, 2 minutes of Random, 3 minutes of long Structured stream, 4 minutes of short Structure blocks, 2 minutes of Random, and 1 minute of RS). The same analysis than above was performed in sliding time windows of 2 minutes with a 1.5 s step.

### 2.6. ERPs to test words

The pre-processed data were filtered between 0.5 and 20 Hz, epoched between [-1.50, 3.25] s from the onset of the triplets. Epochs containing samples identified as artifacts were rejected. On average we retained 24.8 trials for the Word condition (SD 3.6, range [17, 31]), 24.5 for the Edge-word condition (SD 3.6, range [16, 30]), 24.7 for the Part-word condition (SD 4.0, range [16, 30]), and 24.5 for the Non-word condition (SD 4.0, range [14, 30]). Subjects who did not provide at least 12 trials per condition were excluded. Data were reference averaged, normalized by dividing by its standard deviation, and baseline corrected by subtracting the average over the interval between 2.25 s from the onset of the previous word and the corresponding word. Trials were averaged by condition, and two contrasts were studied: *(1) ABx* (Words and Edge-words) vs. *BCx* (Part-words and Non-words) triplets; *(2)* triplets with heard transitions (Words and Part-words) vs. un-heard transitions (Edge-words and Non-words). The responses were compared using non-parametric cluster-based permutation analysis (Oostenveld et al., 2011) in two time windows: *(1)* [0, 0.5] s to detect early effects only attributable to the encoding of the first syllable, and *(2)* [0.5, 2.75] s to detect effects related to a TPs violation or the triplets’ offset. A t-statistic with an alpha threshold of 0.05 was used for clustering; neighbor electrodes had a maximum distance of 3 cm (4.2 neighbors per channel on average); clusters had a minimum size of two, and 5,000 permutations were run to estimate the significance level. The quantification of the effect along test blocks was performed by computing the average difference between *ABx* and *BCx* conditions over the clusters. Data points were included for subjects and blocks when at least 3 out of 8 trials in both conditions were included.

### 2.7. Adult behavioral experiment

33 French speaking adults were tested in an online experiment analogous to the infant study through the Prolific platform and received monetary compensation for their participation. The same stimuli as in the infant experiment were used. Participants first heard 3 minutes of familiarization with the Structured stream. Then, they completed eight sessions of re-familiarization and testing. Each re-familiarization lasted 30 s, and in each test session, all 16 possible test words were presented. Subjects were said they should try to identify the words from an invented language. During the test phase, subjects had to report after each test word their familiarity with it. Subjects responded by using the cursor to click on the screen on a scale from 1 to 6. One participant was excluded because (s)he always responded with a score of 1 or 2. Subjects were randomly assigned to one of the three lists.

## 3. Results

### 3.1. Neural markers of learning in neonates: familiarization phase

During Resting-state, as expected, no entertainment was seen either at the syllabic (4 Hz) or word (1.33 Hz) rates. As expected, for Random streams, we observed enhanced activity at the syllabic rate for many central-frontal and posterior electrodes (*p* < 0.05, FDR corrected) and no enhanced activity at the word rate. During the Structured streams, we observed a similar enhanced oscillatory activity at the syllabic rate and significant neural entrainment at the word rate mainly over left temporal electrodes (*p* < 0.05, FDR corrected) (Figure 2).

**Figure 2.**
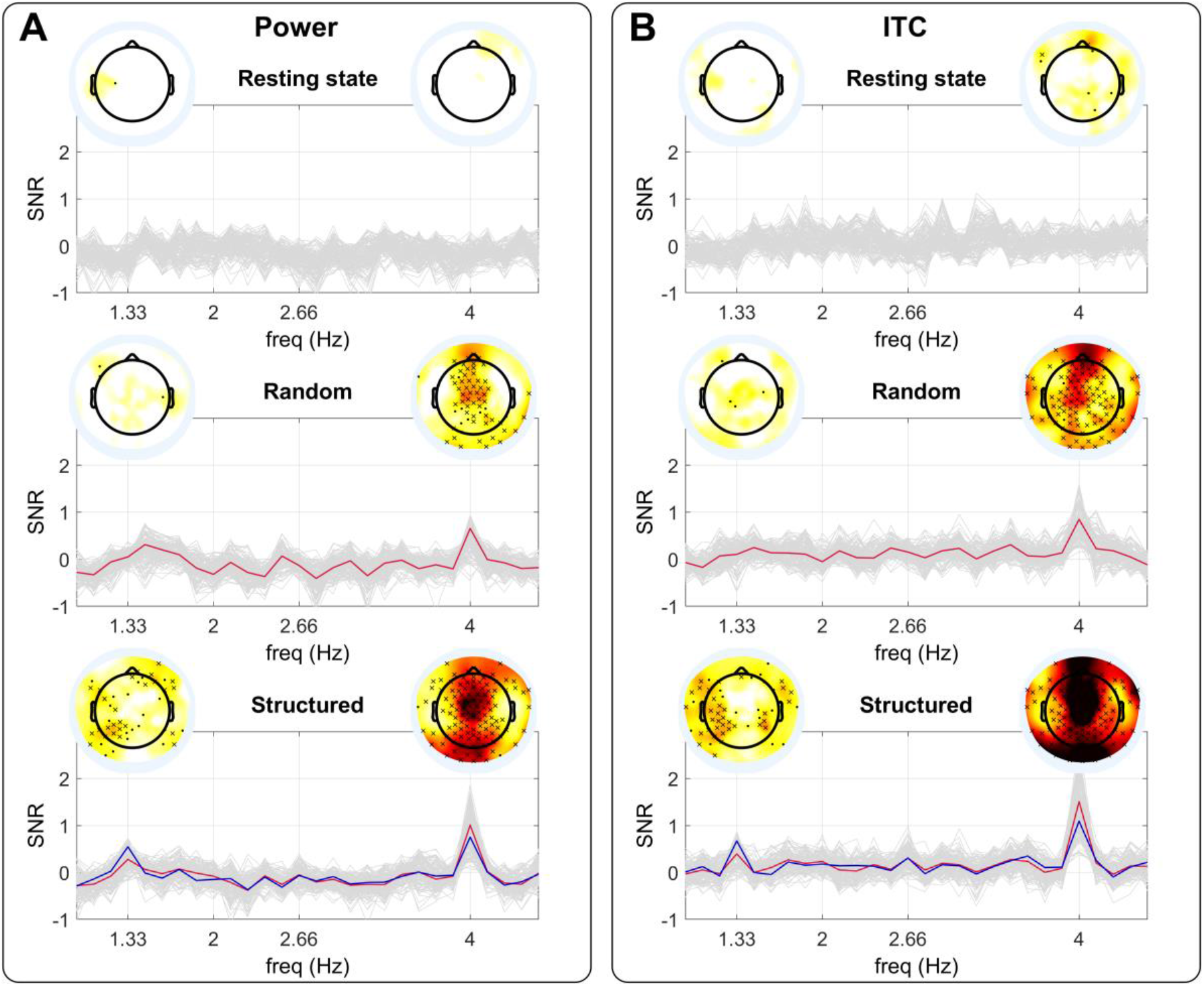
Neural entrainment during Resting-state, Random streams, and Structured streams. **(A)** SNR for the power. In light gray, the entrainment for all electrodes. In red, the mean over the electrodes showing significant entrainment (p < 0.05, one-sided t-test, FDR corrected) at the syllabic rate (4 Hz). In blue, the mean over the electrodes showing significant entrainment (p < 0.05, one-sided t-test, FDR corrected) at the word rate (1.33 Hz). The topographies represent the entrainment in the electrodes space at the word rate (1.33 Hz) and at the syllabic rate (4 Hz). Asterisks indicate the electrodes showing enhanced neural activity (cross: p < 0.05, one-sided t-test, FDR corrected; dot: p < 0.05, one-sided t-test, without FDR correction). **(B)** Same based on ITC.

By comparing the entrainment with a 1-way-ANOVA with condition as within-subjects factor (Fig. 3 A and B), similar results were obtained for power and ITC. A main effect of condition was observed at the syllabic rate (power: *F(2,58) = 21.8, p = 8.6×10^-08^*, ITC: *F(2,58) = 21.8, p = 8.7×10^-8^*, driven by a lower power/ITC during Resting than Random (power:*p = 0.0021*, ITC: *p = 0.0085*) and Structured (*p = 8.4×10^-9^*, ITC: p = 7.5×10^-9^), and lower power/ITC during Random than Structured (power: *p = 0.0075*, ITC: *p = 0.0017*). At word rate there was a main effect of condition (power: *F(2,58) = 10.7, p = 0.00018*, ITC: *F(2,58) = 8.2, p = 0.000706*), due to a higher power/ITC during Structured than Resting (power:*p = 2.9×10^-5^*, ITC: *p = 0.00038*) and Random (power: *p = 0.0052*, ITC:*p = 0.013).* All p-values were Bonferroni corrected for multiple comparisons.

**Figure 3.**
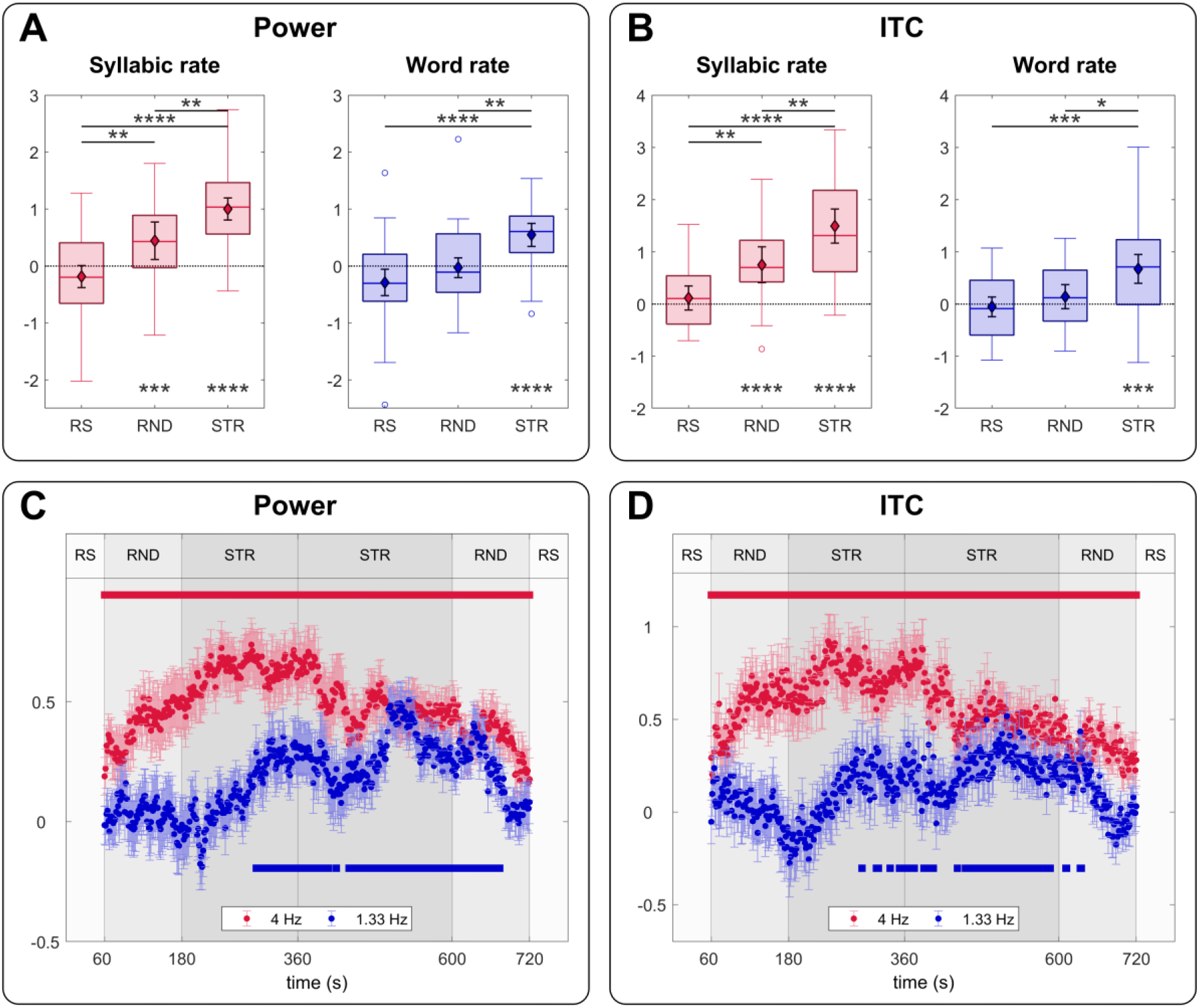
**(A)** SNR for the power at the Syllable and Word rate during the three conditions. Asterisks represent Bonferroni corrected p-values. **(B)** Same as A based on ITC. **(C)** Time course of entertainment based on power computed on 120 s time windows. Error bars represent standard errors. The red line on the top indicates when the power at the Syllabic rate (4 Hz) was bigger than the null hypothesis 0 (p < 0.05, one-sided t-test, corrected by FDR). The blue line on the bottom indicates when the power at the Word rate (1.33 Hz) was bigger than the null hypothesis 0 (p < 0.05, one-sided t-test, corrected by FDR). **(D)** Same as C based on ITC.

To quantify learning through the experiment, we measured entrainment at the syllabic and word rate in sliding time windows of 2 minutes with a 1.5 s step by concatenating the data from all conditions. Notice that because the integration window is two minutes long, the entrainment during the first minute of random, for example includes, data from the structured stream. Results show an increase in Power and ITC at the word rate at around 2 minutes from the beginning of the structured stream (Figure 3 C and D).

### 3.2. Encoding format in neonates: Post-learning phase

We first looked for ERPs components related to ordinal position violations by comparing *ABx* (Words and Edge-words) vs. *BCx* triplets (Part-words and Non-words). A non-parametric cluster-based permutation analysis (Oostenveld et al., 2011) revealed a significant early difference before 500 ms in a positive frontal cluster (*p = 0.0152*, time window [0, 388] ms) and in a left-posterior negative cluster (*p = 0.0324*, time window [0, 308] ms), (Fig. 4 A, B). Given the time window, this effect can only be related to the encoding of the first syllable (i.e., ordinal encoding). A second difference was also observed after the offset of the triplet, in a frontal-left positive cluster (*p = 0.0142*, time window [788, 1600] ms), and even a third one later in a frontal cluster (*p = 0.002*, time window [1684, 2628] ms) (Fig. 4 C, D).

**Figure 4.**
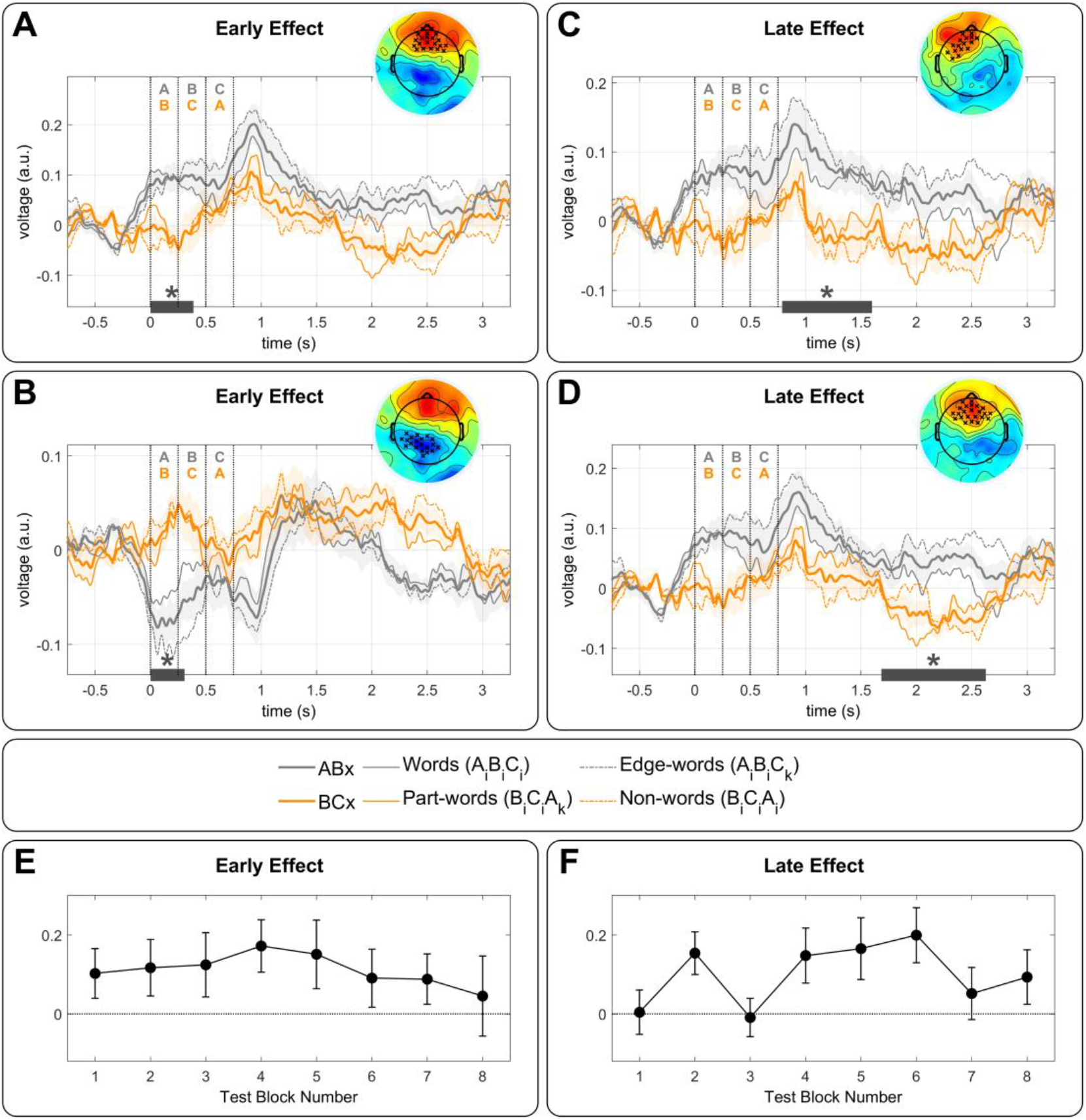
Responses to triplets in isolation. **(A)** Grand-average response to ABx and BCx triplets over the early frontal positive cluster (*p = 0.0152*) obtained from the cluster-based permutation analysis. The thick lines correspond to *ABx* (gray line) and *BCx* (orange line) conditions and the thin lines to the sub-conditions (Words and Edge-Words vs. Part-words and Non-words). Shaded areas correspond to the standard error across neonates. The time zero corresponds to the onset of the test word. The topography shows the difference *ABx-BCx* during the time window where significant differences were observed (gray line under the plot). **(B)** Same as A, but for the early negative cluster over left-temporal posterior electrodes (*p = 0.0324*). **(C)** Same as A, but for a late positive cluster over frontal-left electrodes (*p = 0.0142*). **(D)** Same as A, but for a later positive cluster over frontal electrodes (*p = 0.0020*). **(E)** Time progression of the ERP early effects (A, and B) over the 8 test blocks. **(F)** Time progression of the average of the two late ERP effects (C, and D) over the 8 test blocks.

We then looked for ERPs components related to TPs violations by comparing heard triplets (Words and Part-words) vs. non-heard triplets (Edge-words and Non-words), but we found no significant difference (*p > 0.1*). No significant differences were detected in the comparisons Words vs. Edge-words, and Part-words vs. Non-words (*p > 0.1*).

To ensure that ordinal encoding was present from the beginning of the test phase and was not triggered by hearing isolated triplets, we computed the effect throughout the eight test blocks. Despite fluctuations likely due to the small number of trials, the effect was present from the earliest test blocks (Fig. 4E), suggesting that the encoding of the first syllable in Words had emerged while infants were listening to continuous speech.

### 3.3. Encoding format in adults

Adults rated on a scale their familiarity with the triplets after familiarization with identical streams as neonates (Fig. 5). Results from a linear mixed model using the scoring as dependent variable, the triplet condition as predictor, and subjects as a random factor (*Scoring ~ Cnd + 1 | Sbj*) showed a main effect of condition (*F(3,3721) = 79.72, p < 2.2×10 ^-16^*). A post hoc Tukey test revealed that the Words score was higher than each of the other conditions (*ps < 0.0001*) whereas the Non-words was the lowest, significantly inferior to Part-words (*p < 0.0001*), and to Edge-words (*p = 0.0045*).

**Figure 5.**
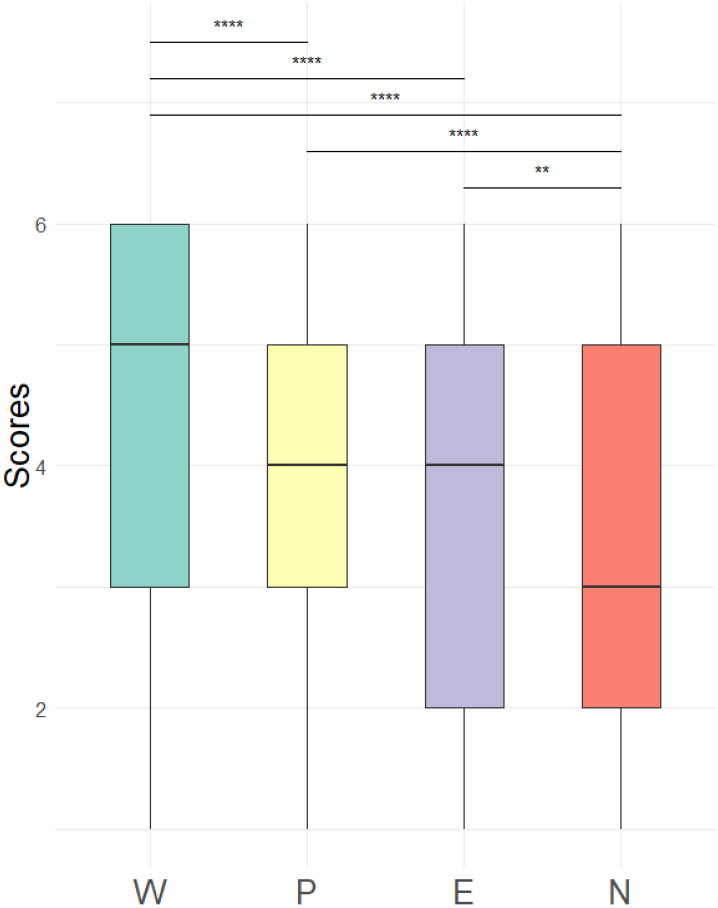
Results for the adults’ behavioral experiment. The distribution of the scores for all trials and participants are represented per condition.

## 4. Discussion

Here, we used a classical speech segmentation task (Saffran et al., 1996) to investigate statistical learning in neonates. While previous studies have shown that infants are sensitive to statistical regularities in speech since birth (Fló et al., 2019; Kudo et al., 2011; Teinonen et al., 2009), it was still unknown what information they tracked and memorized. First, our study revealed that sleeping neonates responded rapidly (within 2 minutes) to the tri-syllabic pattern. Second, when isolated triplets were presented, a differential response was observed from the first syllable, revealing that they expected triplets to start with a specific set of syllables. Third, TP violation did not modulate ERP to triplets. This result indicates a memory representation that no longer depended on TPs, despite TP were used to segment the stream, suggesting a switch to a different format. Finally, these results reveal the accuracy of consonant encoding in newborns that allow them to keep the relationship between 12 syllables and memorize a set of 4 first syllables despite common vowels at different ordinal word positions. This observation is not trivial given the common assumption that infants are initially limited to the most stable units, such as vowels. For example, Benavides et al. (Benavides-Varela et al., 2012) reported a larger novelty response when changing the vowels of a bi-syllabic word (e.g., *lili* to *lala*) compared to a change of consonants (e.g., *lili* to *titi*). However, a recent EEG study showed that phonetic features were at the basis of speech perception in 3-month-old pre-babbling infants, opening the possibility of an innate code based on phonetic features for all phonemes, vowels, and consonants (Gennari et al., 2021).

### 4.1. Learning based on TPs

The significant increase in power and ITC to word rate in the Structured stream demonstrated that TP computations lead to stream structuring. Learning occurred within 2 min of familiarization. This rapid learning is consistent with the length of the stream previously used in behavioral experiments in 8-month-old infants (Saffran et al., 1996) and EEG experiments in adults and 6-month-old infants (Batterink & Choi, 2021; Benjamin et al., 2021). The concordance of learning rate across ages indicates that statistical learning abilities do not improve markedly with age, a remarkable observation given the significant maturational changes in auditory/linguistic regions and hippocampus during the first years of life (Adibpour et al., 2020; Leroy et al., 2011).

We did not characterize the neonates’ sleep stages. However, their general behavior during the recording session (eyes closed, hypotonia), the duration of the experiment, and the lack of task and reward, combined with the short awake periods outside of feeding in the days after birth, certainly did not favor an attentive and focused listening of the auditory input. Neonates’ success in extracting the regularities is congruent with adult studies showing neural entrainment at the word rate even when participants are distracted by a primary task (Batterink & Choi, 2021; Benjamin et al., 2021), revealing the automaticity of TP calculations.

In adult experiments, the word rate entrainment is accompanied by decreased syllabic rate entrainment (Batterink & Choi, 2021). Our results revealed a more complex pattern. The syllabic rate entrainment increased at the beginning of the Structured stream and decreased when word rate entrainment became significant. The initial increase entrainment at the syllabic rate might reflect stronger activation of the language network during the uncovering of the structure compared to random syllable presentation. This hypothesis would be consistent with an adult functional magnetic resonance imaging (fMRI) experiment showing that activity in the left-temporal cortex is modulated by the level of complexity of speech sequences (Pallier et al., 2011). The subsequent decrease might result from top-down inhibition of the syllabic response once the stream has been segmented.

While neural entrainment demonstrated that infants were sensitive to the rhythmic structure of the stream, this might result from an automatic error response elicited by the unpredictability of the first syllable (TPs) or by a neural response to tri-syllabic chunks (segmentation). The responses to isolated triplets dissociated these two alternatives.

### 4.2. Memory representation of the segmented words

ERPs to the isolated triplets revealed the format of the memorized information. ERPs differed from the first syllable between *ABx* triplets (Words and Edge-Words) and *BCx* triplets (Parts-Words and Non-Words); thus, before any TP violation (*AB* and *BC* transitions were both equal to 1). Additionally, we observed no specific ERP component after a TPs violation, that is to say, between Words and Edge-Words on one side and Part-Words and Non-Words on the other side. It is important to note that in Non-words, the first syllable was presented at the last position without evoking a particular response (i.e., a difference with Part-Words). The absence of a distinctive response to the first syllable at the wrong position favors the hypothesis that it is not a particular familiarity with this syllable that caused the difference between *ABx* and *BCx* triplets, but its position as a first syllable in the triplet.

Two approaches have been proposed for flat continuous speech segmentation. From one perspective, the TPs are computed, and the drops in TPs serve as cues to word boundaries (Saffran et al., 1996). From another perspective, recurrent chunks of co-occurring syllables are identified and stored in memory (Perruchet & Vinter, 1998). Our experiment did not attempt to disentangle these two mechanisms. However, the lack of difference between heard and un-heard triplets revealed that neonates retained neither the full TP matrix nor the entire Words. Instead, they remained limited to the memorization of the first syllable. Meanwhile, adults scored Words as highly familiar, Edge-words as more familiar than Non-words, and finally Edge-Words and Part-words as equally familiar (although Edge-words never appeared in the stream). These results suggest that adults memorized the complete Words, but also that the format of the representation depended on both TPs and ordinal encoding, in agreement with other recent studies (Fló, 2021; Henin et al., 2021).

Altogether, our results suggest a multistep process. Segmentation occurred either because the drop in TP produced a prediction error that singularized the non-predicted syllable (i.e., the first syllables) or because the association of the syllables within words increased their similarity, leading to boundaries at the landscape lower points. In a second step, the segmented triplets are stored in memory, and therefore, any memory constraints might affect the retained representation.

### 4.3. Word memorization is incomplete in neonates

Neonates are thus memorizing at least the first syllable of the chunk pointing to an ordinal encoding, the third level of complexity in Dehaene et al. taxonomy (Dehaene et al., 2015). However, they did not distinguish Words (*A_i_B_i_C_i_*) and Edge-words (*A_i_B_i_C_k_*), suggesting that neonates’ words memory was not complete. A limited memory capacity in neonates for middle positions has already been described. A NIRS study showed a better encoding of the syllables at the edges of a six-syllable pseudo word than in intermediate positions (Ferry et al., 2016). Unfortunately, the conditions in that study do not allow disentangling if the effect was due to a better encoding of the first, the last, or both syllables. A memory capacity limited to the first syllable might explain previous neonates studies such as the recognition of bi-syllabic pseudo-words from a new pseudo-word presented two minutes later (Benavides-Varela et al., 2012; Benavides-Varela & Mehler, 2011) and of words conforming a structured stream (Fló et al., 2019).

This memory limitation might be due either to immaturity or to sleep. Sleep is primarily considered as consolidating memories, and while learning is suppressed during deep non-REM stage in adults, implicit word learning is present during REM sleep (Andrillon et al., 2017). Thus, sleep might not represent the limiting factor here, and this question should be further studied.

### 4.4. Putative underlying neural networks

While EEG has an excellent temporal resolution, it does not provide accurate spatial resolution and information regarding the activity of brain structures. However, we may speculate from the adults’ results and the few brain imaging studies in infants investigating the maturation of the pertinent brain regions. Henin and colleagues (Henin et al., 2021) isolated three main networks in a similar task in epileptic patients that might already be at work in neonates. The superior temporal region, which might be related to local processes involved in TP computations, and two memory structures: the dorsal linguistic pathway supporting verbal working memory, and the hippocampus, recently reported as engaged in sequence learning (Schapiro et al., 2014; Schlichting et al., 2017). Although these two structures have been considered immature in infants, fMRI has revealed that they support cognitive functions in the first trimester. Notably, whereas the superior temporal regions are affected by the immediate repetition of a sentence (Dehaene-Lambertz, 2017), repetition at a longer time-scale of 14 seconds produces activation in the inferior frontal gyrus in three-month-old infants (Dehaene-Lambertz et al., 2006). Moreover, a NIRS study in sleeping neonates revealed that a correlated activity between left-temporal and left-frontal regions, compatible with activation in the dorsal linguistic pathway, is crucial for word learning (Benavides-Varela et al., 2017). As for the hippocampus, activity has been reported in infants as young as 3-months when performing a visual sequence learning task, with no modulation by infant’s age (Ellis et al., 2020). Thus, future work should investigate whether hippocampal circuits considered fundamental to SL, such as the monosynaptic pathway, are involved in such a word-learning task since birth. fMRI in infants might help determine how the network highlighted in adults (Henin et al., 2021) is similarly involved in infants to support the two stages we have isolated the relative role of the hippocampus and the linguistic network.

## 5. Conclusions

Despite their unquestionable immaturity, neonates reveal sophisticated learning abilities. From drops in TPs, they were able to segment a continuous speech stream and start to encode the first syllables of the chunks. While the present study remains a toy experiment far from the complexity of a real-life environment, it reveals the necessary integration between successive functional processes computed in different neural structures that is at the core of learning in infants.

## Acknowledgments

This research has received funding from the European Research Council (ERC) under the European Union’s Horizon 2020 research and innovation program (grant agreement No. 695710).

## Author Contributions

A.F. and G.D.L. designed the research; A.F., L.B. and M.P. performed the research; A.F. analyzed the data; and A.F., L.B., and G.D.-L. wrote the paper

